# Genotypic and phenotypic characterization of *Enterococcus faecalis* isolates from periprosthetic joint infections

**DOI:** 10.1101/2024.02.06.579140

**Authors:** Amanda Haeberle, Kerryl Greenwood-Quaintance, Sarah Zar, Stephen Johnson, Robin Patel, Julia L. E. Willett

## Abstract

Over 2.5 million prosthetic joint implantation surgeries occur annually in the United States. Periprosthetic joint infections (PJIs), though occurring in only 1-2% of patients receiving replacement joints, are challenging to diagnose and treat and are associated with significant morbidity. The Gram-positive bacterium *Enterococcus faecalis*, which can be highly antibiotic resistant and is a robust biofilm producer on indwelling medical devices, accounts for 2-11% of PJIs. *E. faecalis* PJIs are understudied compared to those caused by other pathogens, such as *Staphylococcus aureus*. This motivates the need to generate a comprehensive understanding of *E. faecalis* PJIs to guide future treatments for these infections. To address this, we describe a panel of *E. faecalis* strains isolated from the surface of prosthetic joints in a cohort of individuals treated at Mayo Clinic in Rochester, MN. Here, we present the first complete genome assemblage of *E. faecalis* PJI isolates. Comparative genomics shows differences in genome size, virulence factors, antimicrobial resistance genes, plasmids, and prophages, underscoring the genetic diversity of these strains. These isolates have strain-specific differences in *in vitro* biofilm biomass, biofilm burden, and biofilm morphology. We measured robust changes in biofilm architecture and aggregation for all isolates when grown in simulated synovial fluid (SSF). Lastly, we evaluated antibiotic efficacy of these isolates and found strain specific changes across all strains when grown in SSF. Results of this study highlight the existence of genetic and phenotypic heterogeneity among *E. faecalis* PJI isolates which will provide valuable insight and resources for future *E. faecalis* PJI research.

**Importance:** Periprosthetic joint infections (PJIs) affect ∼1-2% of those who undergo joint replacement surgery. *Enterococcus faecalis* is a Gram-positive opportunistic pathogen that causes ∼10% of PJIs in the United States each year, but our understanding of how and why *E. faecalis* causes PJIs is limited. *E. faecalis* infections are typically biofilm associated and can be difficult to clear with antibiotic therapy. Here, we provide complete genomes for four *E. faecalis* PJI isolates from the Mayo Clinic. These isolates have strain-specific differences in biofilm formation, aggregation, and antibiotic susceptibility in simulated synovial fluid. These results provide important insight into genomic and phenotypic features of *E. faecalis* isolates from PJI.

## Introduction

*Enterococcus faecalis* is a Gram-positive, facultative anaerobic bacteria that colonizes the human gastrointestinal tract and remains a minor component of the healthy microbiota in adults. It is also a prolific opportunistic pathogen that causes biofilm-associated infections such as urinary tract infections, infected root canals, infective endocarditis, and periprosthetic joint infections (PJI), and is a leading cause of hospital acquired infections^1–6^. Its pathogenicity is due to its ability to form biofilms, which can complicate treatment as biofilms are difficult to remove from tissue and indwelling medical devices. Additionally, biofilms increase the intrinsic antibiotic resistance of *E. faecalis*^7,8^.

PJIs are the result of either perioperative contamination or spread of bacteria from a distant site of infection to the joint, either continuously or hematogenously^9^, and are characterized by inflammation of the joint region with symptoms including pain, joint swelling and/or effusion. These infections are difficult to treat, requiring surgical intervention and prolonged antibiotic treatment^10,11^. Current antibiotic treatment for *E. faecalis* PJIs is controversial and consists mainly of a cocktail of aminopenicillin derivatives, vancomycin, linezolid, and aminoglycosides^12^. *E. faecalis* accounts for 2-11% of all PJIs and is understudied as compared to *Staphylococcus aureus* and *Staphylococcus epidermidis*, the most common PJI pathogens^5,13,14^. While *E. faecalis* is less isolated from PJIs compared to *S. aureus* and *S. epidermidis, E. faecalis* PJIs have a high rate of treatment failure and morbidity for patients, thus representing an area of unmet need in PJI research^5,12^.

The purpose of this study was to gain insight into *E. faecalis* PJIs by characterizing the genomic and phenotypic features of 4 *E. faecalis* isolates from PJIs from patients treated at the Mayo Clinic in Rochester, Minnesota. We generated the first fully assembled genomes of *E. faecalis* PJI isolates and analyzed plasmids, antibiotic resistance genes, virulence factors, core and accessory genes, and prophages. We evaluated their ability to grow, form biofilms and auto aggregate in standard growth media and in simulated synovial fluid (SSF) and found vast differences in these phenotypes when these isolates were grown in SSF. Furthermore, we found differences in the susceptibility of each isolate to antibiotics commonly used to treat enterococcal infections after growth in SSF. These findings underscore the need to use *in vitro* conditions that model relevant host conditions and provide a platform for pursuing future functional genomic and pathogenesis studies of *E. faecalis* PJIs.

## Results

### Clinical presentation, patient management, and identification of *E. faecalis* in PJIs

Subjects were referred to Mayo Clinic for infection and underwent two-stage resection and revision with an antimicrobial cement spacer placed and antimicrobial treatment prior to reimplantation. *E. faecalis* was cultured from sonicate fluid (numbered one through five by date of resection surgery) from five subjects classified as having PJI caused by *E. faecalis* between March 2005 and September 2009. Subject age and gender, implant age at resection, preoperative antimicrobial usage, and the identification number of the corresponding *E. faecalis* isolates that were grown from culture of the sonicate fluid are shown in **Table 1**. All subjects were doing well at their last follow-up.

**Table 1.** Patient information.

| Sonicate Fluid Sample # | Subject Age at Implant Resection (years) | Gender | Implant Location | Implant Age at Resection Surgery (months) | Preoperative Antimicrobial Usage; Timeline of Discontinuation | Preoperative *Enterococcus faecalis* Cultured Strain Identification |
| --- | --- | --- | --- | --- | --- | --- |
| 1 | 80 | Male | Hip | 13 | Vancomycin, rifampin, trimethoprim/sulfamethoxazole; stopped 2 weeks before surgery | IDRL-7415 |
| 2 | 44 | Female | Hip | 22 | Ceftriaxone, moxifloxacin; stopped 2 weeks before surgery | IDRL-7538 |
| 3 | 83 | Male | Knee | 12 | Rifampin, trovafloxacin; stopped 1 week before surgery | IDRL-7639 |
| 4 | 59 | Male | Knee | 19 | Ampicillin; stopped 2 weeks before surgery | IDRL-9065 |
| 5 | 80 | Female | Hip | 32 | None | IDRL-8618 |

### Genomics of *Enterococcus faecalis* isolates from periprosthetic joint infections

*E. faecalis* remains a difficult pathogen to treat during a PJI and little work has been done to understand *E. faecalis* pathogenesis in PJI. To date, no fully assembled genomes of *E. faecalis* isolated from PJIs exist, making it difficult to pursue relevant genomic and mechanistic investigations on *E. faecalis* adaptations in this host region. To address this knowledge gap, we used short-read Illumina and long-read Nanopore sequencing for hybrid genome assembly of each isolate. Bandage plots were used to evaluate closure of each genome (**Supplementary** Figure 1)^15^. Genomes were assembled in Unicycler, annotated using RASTtk, and subject to the Comprehensive Genome Analysis platform in BV-BRC for further investigation. We were unable to generate a closed genome for IRDL-8618, and we observed multiple colony morphologies for this isolate when grown in rich medium. All other isolates had one morphology when grown on rich medium. Therefore, we pursued further investigation for the four remaining isolates (IDRL-7415, IDRL-7538, IDRL-7639, and IDRL-9065) (**Figure 1**). These are the first fully assembled genomes for *E. faecalis* PJI isolates making them a valuable resource for future genomic and phenotypic studies.

**Figure 1.**
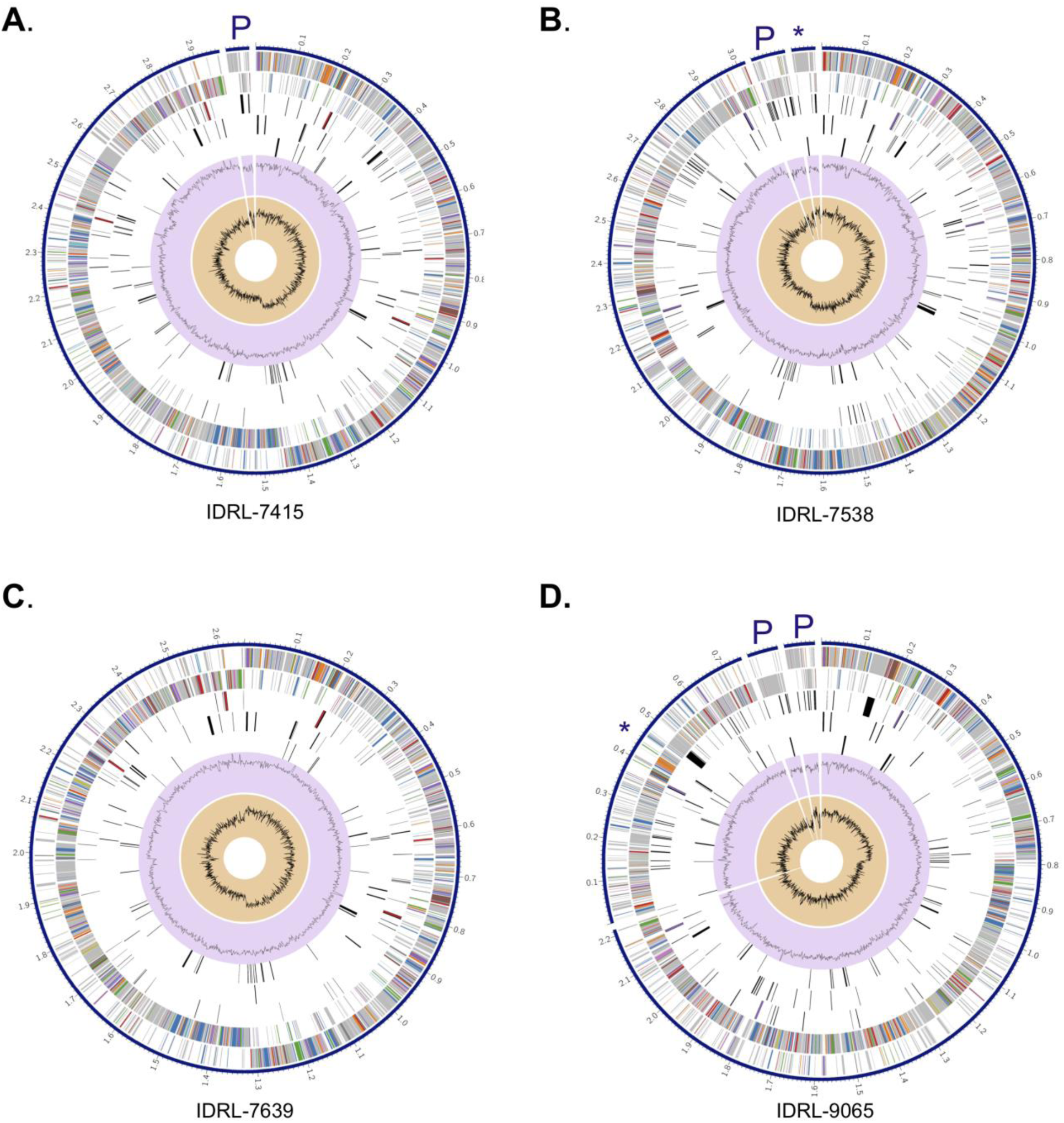
Fully Assembled Genomes of PJI isolates of *E. faecalis*. Circular graphical display of the distribution of each genome assembled with short and long read sequencing in Unicycler and annotated with RAStk for **(A)** IDRL-7415, **(B)** IDRL-7538, **(C)** IDRL-7639, and **(D)** IDRL-9065. Rings show (from outer to inner) the contigs, CDS forward strand, CDS reverse strand, RNA genes, CDS homologous to known antimicrobial resistance genes, CDS with homology to known virulence factors, GC content and GC skew. Asterisks (*) indicates the location of genes related to pheromone-inducible plasmids, and P indicates additional plasmids identified using BV-BRC and/or PlasmidFinder.

Genome sizes varied between isolates, ranging from 2.65 Mb to 3.16 Mb (**Table 2**). Similarly, each isolate contained differing functionally assigned and hypothetical proteins, antimicrobial resistance genes, and virulence factors (**Table 2**), a common observation in clinical isolates of *E. faecalis*^16^. Furthermore, there were numerous predicted plasmids in three of four isolates. These included a pheromone-inducible conjugative plasmid, which has a role in cell-cell aggregation and DNA transfer,^17^ in IDRL-7538 and IDRL-9065. Five of the six plasmids predicted by BV-BRC in IDRL-7415, IDRL-7538 and IDRL-9065 were also identified using PlasmidFinder^18^ (**Figure 1, Supplementary Table 1**). We compared the isolates to each other by constructing a bacterial genome tree using RAxML^22^ (**Figure 2A**). We also included the well-characterized strain *E. faecalis* OG1RF as it has well-established genetic tools and has been used for numerous infection and biofilm studies^19–21^. IDRL-7415 was most closely related to OG1RF, followed by IDRL-7538. IDRL-9065 and IDRL-7639 were more closely related to each other than to OG1RF (**Figure 2A**). These findings highlight genetic differences between these PJI isolates and suggest the importance of using them for further *E. faecalis* PJI investigations.

**Figure 2.**
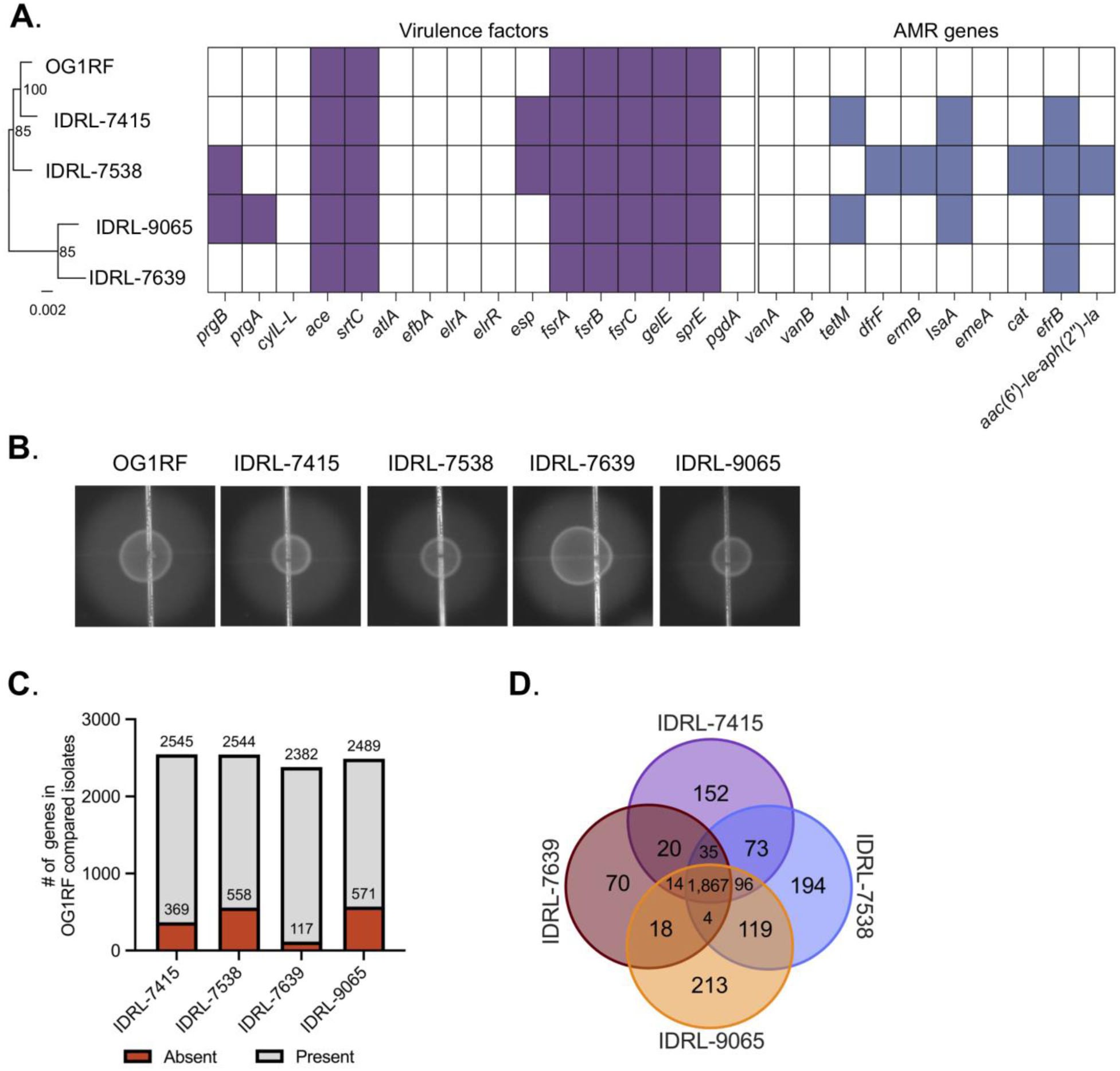
Comparative genomics reveal varying genomic characteristics of PJI isolates. **(A)** Phylogenetic tree generated with the Codon Tree method in BV-BRC. Bootstrap values are indicated at the nodes (100 rounds of “rapid” bootstrapping in RAxML^37^). The text of each node label was updated in the final figure. Shaded regions in the table indicate the presence of virulence factors (purple) or antibiotic resistance genes (blue). **(B)** Validation of GelE activity in each isolate via gelatinase assay on agar plates supplemented with 3% gelatin. **(C)** Distribution of the number of present and absent genes in OG1RF as compared to each PJI isolate. The area shaded red indicates the number of genes in each PJI isolate that are not found in OG1RF. **(D)** Comparison of core and accessory protein families among OG1RF and PJI isolates (2930 total protein families identified).

**Table 2.** Summary of *E. faecalis* genome findings.

| Isolate | Genome length (bp) | # of contigs | Plasmids | # of hypothetical proteins/ functionally assigned proteins | # of antibiotic resistance genes (PATRIC) | # of virulence factors (VFDB) |
| --- | --- | --- | --- | --- | --- | --- |
| IDRL-7415 | 3,012,124 | 2 | 1 | 613/2,299 | 34 | 20 |
| IDRL-7538 | 3,161,211 | 3 | 2 | 693/2,409 | 38 | 30 |
| IDRL-7639 | 2,652,177 | 1 | 0 | 445/2,054 | 32 | 16 |
| IDRL-9065 | 3,105,522 | 4 | 3 | 707/2,353 | 34 | 23 |

We screened each *E. faecalis* PJI isolate genome and OG1RF for virulence factors using the Comprehensive Genome Analysis tool in BV-BRC^22^. All four isolates contained genes encoding Ace, a collagen adhesion protein involved in pathogenesis of *E. faecalis*^23^, and Ebp pili^24^. Additionally, all four isolates had genes encoding the Fsr quorum sensing system and the quorum sensing-controlled proteases SprE and GelE. GelE is a metalloprotease that mediates chain length and autolysis, host intestinal epithelial permeability, and biofilm formation^25–28^ (**Figure 2A**). Because discrepancies between gelatinase genotype and phenotype have previously been described in clinical isolates^29,30^, we determined if functional GelE was produced by performing gelatinase assays for all four strains. The GelE-positive strain OG1RF was included as a control. All four PJI isolates produced gelatinase, as indicated by the development of a halo around colonies grown on an agar plate containing gelatin (**Figure 2B**).

OG1RF is a strain that is commonly used for functional genomic studies^19–21^. However, this strain was originally isolated from the oral cavity, suggesting that its genetic adaptations may be different from that of other clinical isolates of *E. faecalis.* Therefore, we wanted to compare the genome of OG1RF to each PJI isolate to determine whether they contain genes that are not found in the OG1RF background. Each isolate shared ∼2,300 genes with OG1RF (**Figure 2C**). Additionally, IDRL-7639 had 117 genes not found in OG1RF, followed by IDRL-7415 (369 genes), IDRL-7538 (558 genes), and IDRL-9065 (571 genes). We next determined the core and unique protein families in each isolate. All four isolates shared a core set of 1,867 protein families with variation among the number of unique protein families. IDRL-7639 had 70 unique protein families not found in other clinical isolates, followed by DRL-7415 (152 protein families), IDRL-7538 (194 protein families), and IDRL-9065 (213 protein families).

We also screened each *E. faecalis* PJI isolate genome for antimicrobial resistance genes and prophages. All four isolates were predicted to encode the ABC transporter EfrBCD involved in mediating multidrug efflux in *E. faecalis* **(Figure 2A)**^31^. Three of the four isolates had the gene *lsaA*, which encodes another efflux pump that contributes to resistance to clindamycin and quinupristin-dalfopristin^32^ **(Figure 2A)**. To further evaluate the antimicrobial resistance of these clinical isolates, we carried out minimum inhibitory concentration (MIC) assays on all four strains. Two colony morphologies were observed for IDRL-9065 when growing isolates for these experiments, so a representative colony of each size was used for MIC assays. Only one strain (IDRL-7538) displayed resistance to an antimicrobial according to clinical breakpoints (**Table 3**). This strain harbors the *aac(6’)-Ie-aph(2”)-Ia* gene, which encodes an aminoglycoside-modifying enzyme that confers high-level gentamicin resistance, and the isolate had gentamicin synergy resistance in the MIC assay (**Figure 2A**, **Table 3**)^33^. We also found strain-specific differences in the presence of prophages using PHASTER^34^ (**Supplementary** Figure 2). Each isolate except for IDRL-7639 had complete prophage sequences encoded on their genome. phiEf11, which was originally identified as a temperate phage in an oral isolate of *E. faecalis*^35^, was found in both IDRL-7415 and IDRL-9065. phiFL4A, originally identified as a temperate phage in a bacteremia isolate^36^, was found in both IDRL-7538 and IDRL-9065 (**Table 3**). Taken together, this analysis shows genomic diversity in virulence factors, antibiotic resistance genes and prophages in these *E. faecalis* PJI isolates.

**Table 3.**
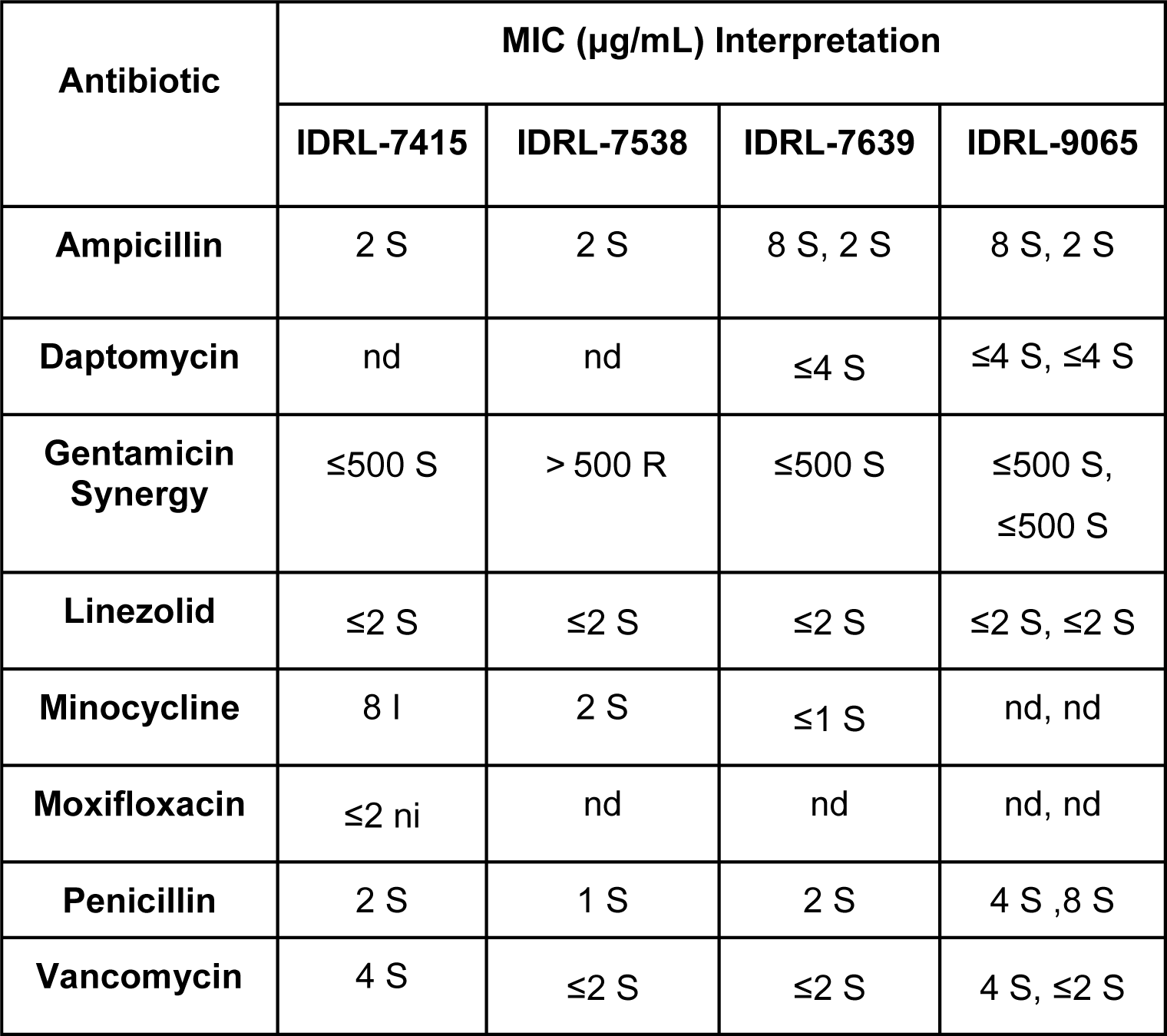
MIC analysis of PJI isolates. Antimicrobial susceptibility interpretation was determined according to Clinical and Laboratory Standards Institute guidelines. MIC, minimal inhibitory concentration; S, susceptible; I, intermediate; R, resistant; ni, no interpretive guideline; nd, not done. IDRL-9065 values were reported for 2 colony morphologies.

### *E. faecalis* PJI isolates have variable biofilm morphology, which is altered in simulated **synovial fluid**

Given the genetic diversity of these isolates, we wondered whether biofilms produced by these strains would be similar or whether they would display variation in characteristics such as biofilm biomass accumulation or biofilm architecture. OG1RF is one of the most commonly used strains to study *E. faecalis* biofilms and typically forms flat biofilms *in vitro* with some small chain-like structures^19^. We hypothesized that differences in genome composition of PJI isolates would contribute to diverse *E. faecalis* biofilm properties. To evaluate biofilm growth, we visualized biofilms using fluorescence microscopy after 6 h (early) and 24 h (late) growth in BHI (**Figure 3A**). Morphology differences between isolates were evident at both time points. Similar to previous reports, OG1RF biofilms consisted of individual diplococci and short chains. IDRL-7415 and IDRL-7639 had comparable biofilm morphologies to OG1RF. Conversely, IDRL-7538 and IDRL-9065 consisted of some individual cells with longer chains of cells present (**Figure 3A**). Furthermore, IDRL-6538, IDRL-7639, IDRL-9065 had greater cell densities at 6 h as compared to 24 h. Conversely, OG1RF and IDRL-7415 had greater cell densities at 24 h (**Figure 3A**).

**Figure 3.**
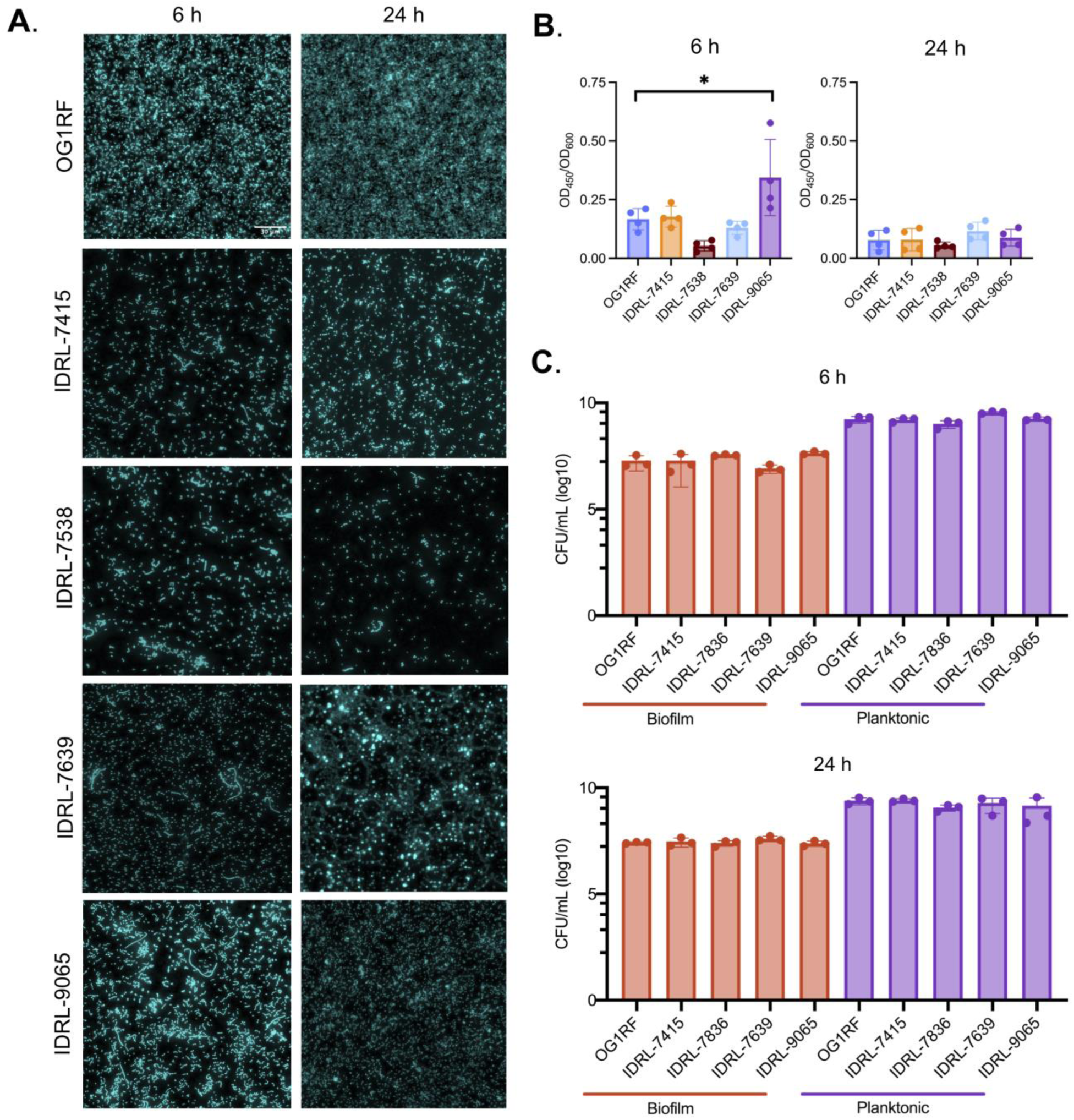
Phenotypic characteristics of PJI isolates. Each isolate was subjected to *in vitro* assays in BHI to assess their **(A)** biofilm architecture, **(B)** biofilm production in 96-well plates, and **(C)** planktonic and biofilm cell viability using a submerged substrate assay. All assays were done at 6 and 24 h post-inoculation. Data represents n = 3 independent biological replicates for panels **(A)** and **(C)**, and n = 4 independent biological replicates for panel **(B)**. Statistical significance was determined for **(B)** and **(C)** using ordinary one-way ANOVA with Dunnett’s test for multiple comparisons. ns = P > 0.05, * = P ≤ 0.05. The scale bar (shown in OG1RF) represents 30 μm.

We next measured biofilm biomass accumulation for all four isolates using microtiter plate biofilm assays. Previous studies of *E. faecalis* biofilms have demonstrated that different assays can provide different measurements of biofilm production^16,19^. Safranin staining in microtiter plate biofilm assays can measure biofilm biomass, compromising bacterial cells and extracellular polymeric substances^38^. We measured total bacterial cell growth (OD_600_) and biofilm biomass (OD_450_) after 6 and 24 h growth in BHI. At 6 h, we observed more variability in the biofilm biomass to cell growth ratio (biofilm index OD_450_/OD_600_) between isolates compared to 24 h. IDRL-7415 and IDRL-7639 were similar to OG1RF. IDRL-7538 had the lowest biofilm index while IDRL-9065 had the highest (**Figure 3B**). At 24 h, the biofilm index decreased for all strains relative to 6 h (**Figure 3B**). Finally, we measured biofilm burdens for each isolate using submerged substrate assays, in which strains were cultured in multiwell plates containing Aclar discs. This approach allows for isolation and enumeration of both viable planktonic and biofilm burden via CFU quantification^19,39^. At both 6 h and 24 h, the biofilm CFU/mL burden for each isolate was similar to that of OG1RF (∼10^7^ CFU/mL) (**Figure 3C**). Interestingly, IDRL-9065 had the highest biofilm index at 6 h (as measured by safranin staining) but similar biofilm burden (as measured by CFU/mL). Together, these results demonstrate heterogeneity in biofilm growth across *E. faecali*s PJI clinical isolates.

### Simulated synovial fluid induces changes in *E. faecalis* growth and biofilm formation

Previous studies have demonstrated that synovial joint fluid induces dramatic changes in *Staphylococcus* growth and biofilm structure *in vivo* and *in vitro* relative to standard growth media^40–43^. Therefore, to reflect the *in vivo* environment for studying *E. faecalis* PJI pathogenesis, we used simulated synovial fluid (SSF) to assess whether *E. faecalis* growth and biofilm formation was different in SSF compared to standard growth conditions. Growth in SSF was evaluated by titrating SSF into BHI and measuring the OD_600_ of OG1RF over time (**Supplementary** Figure 3). Overall, OG1RF grew less in SSF. For additional studies, we chose to use 70% BHI + 30% SSF as this still supported OG1RF growth. Growth in this condition was similar for all four PJI clinical isolates (**Supplementary** Figure 4).

Next, we examined biofilm architectures grown in 70% BHI + 30% H_2_O (control) and 70% BHI + 30% SSF. Because we observed the most variability in biofilm formation between strains at 6 h (**Figure 3B**), we chose this time point for biofilm studies with SSF. Cultures were incubated for 6 h in optically clear microtiter plates, and attached biofilm was stained with Hoechst 33372. We used fluorescence microscopy to visualize cellular organization. Strikingly, unlike biofilms grown in control conditions, larger clumps or aggregates were evident in SSF for three of the four clinical isolates. (**Figure 4A**). Overall, OG1RF and IDRL-9065 biofilms had similar small clumps, and IDRL-9065 grew in long chains. IDRL-7538 and IDRL-7639 had observable aggregates in SSF but not under control conditions. These observations demonstrate that biofilms made by different *E. faecalis* isolates have different morphologies. Next, we measured biofilm biomass accumulation in SSF for all four isolates using microtiter plate biofilm assays and safranin staining as described above. Each isolate was inoculated in control and SSF conditions. There was no significant difference in biofilm index for any strain in SSF compared to control conditions. Together, these data suggest that SSF does not affect biofilm biomass accumulation as measured by safranin staining in microtiter plates but does influence biofilm structure. Future investigations will be necessary to better understand the mechanisms underlying *E. faecalis* biofilm morphology and survival in SSF.

**Figure 4.**
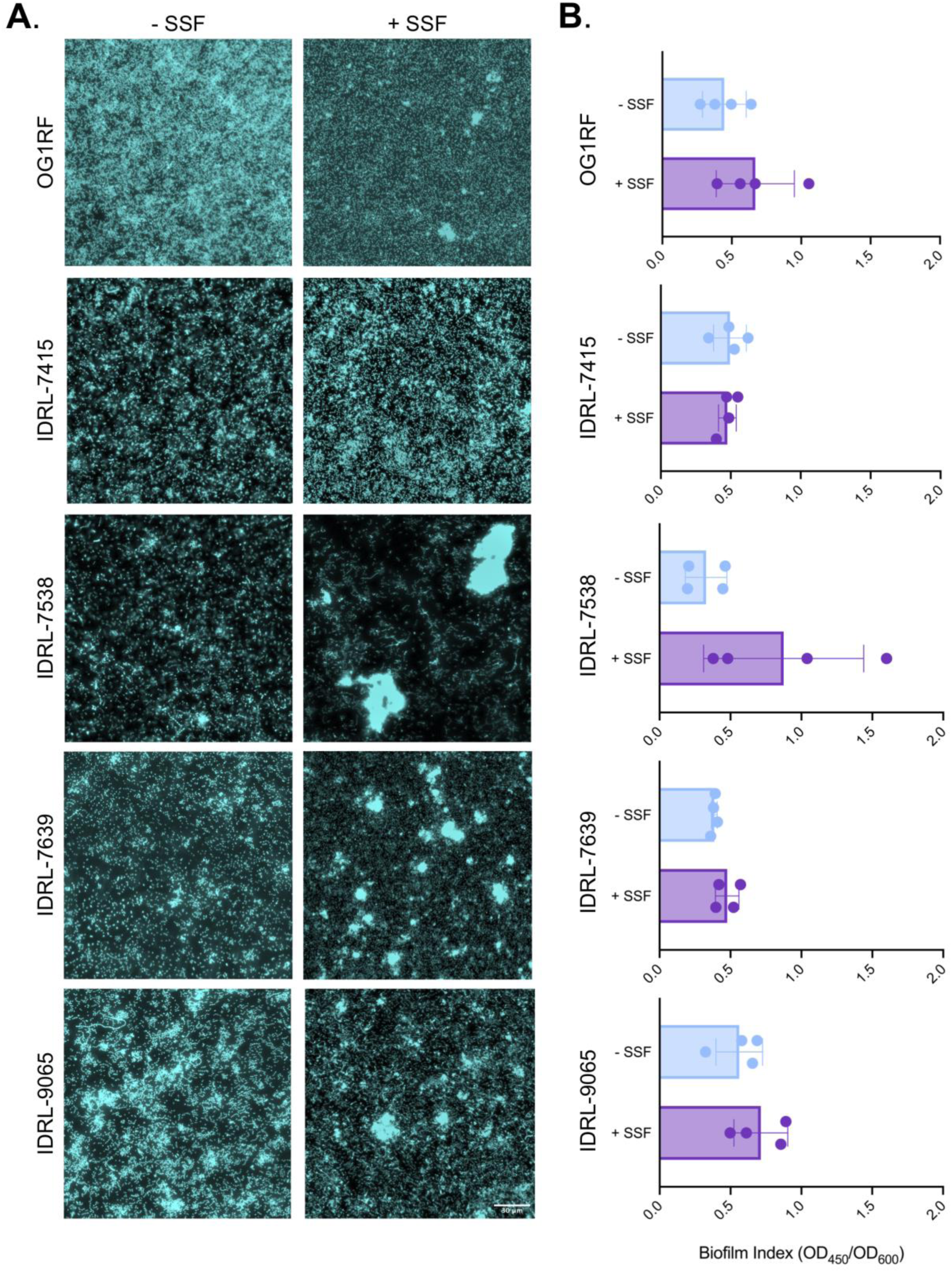
*E. faecalis* biofilm architecture is altered by growth in SSF. **(A)** Biofilm formation of each isolate was observed at 6 h using fluorescence microscopy. Hoechst staining and fluorescence microscopy demonstrated unique architectural differences among clinical isolates grown in BHI vs SSF. Data represents n = 3 independent biological replicates. Scale bar (shown on IDRL-9065) represents 30 μm. **(B)** 96-well plate plate biofilm assay of each isolate grown for 6 h in control and SSF conditions. Data represents n = 4 biological replicates.

### SSF induces autoaggregation of *E. faecalis* planktonic cultures

SSF induces aggregation in both *S. aureus* and *S. epidermidis*^40,42^. Aggregation of bacterial cells also promotes antimicrobial resistance during infection, which can complicate treatment of PJIs^44^. Therefore, we sought to determine if SSF promotes aggregation in *E. faecalis* PJI isolates. We quantified aggregation of overnight cultures using OD_600_ values measured before and after mixing culture tubes (**Figure 6A**). Interestingly, every *E. faecalis* PJI clinical isolate and OG1RF had a significant increase in aggregation in SSF as compared to control growth conditions (**Figure 6**). Aggregation was visible at the bottom of test tubes grown in SSF conditions, but not in BHI controls. OG1RF and IDRL-7538 had ∼3-fold higher aggregation in SSF as compared to BHI controls (**Figure 6BD**, respectively). IDRL-7415, IDRL-7639, and IDRL-9065 had statistically significant increases in aggregation compared to BHI controls (**Figure 6CEF**, respectively).

**Figure 6.**
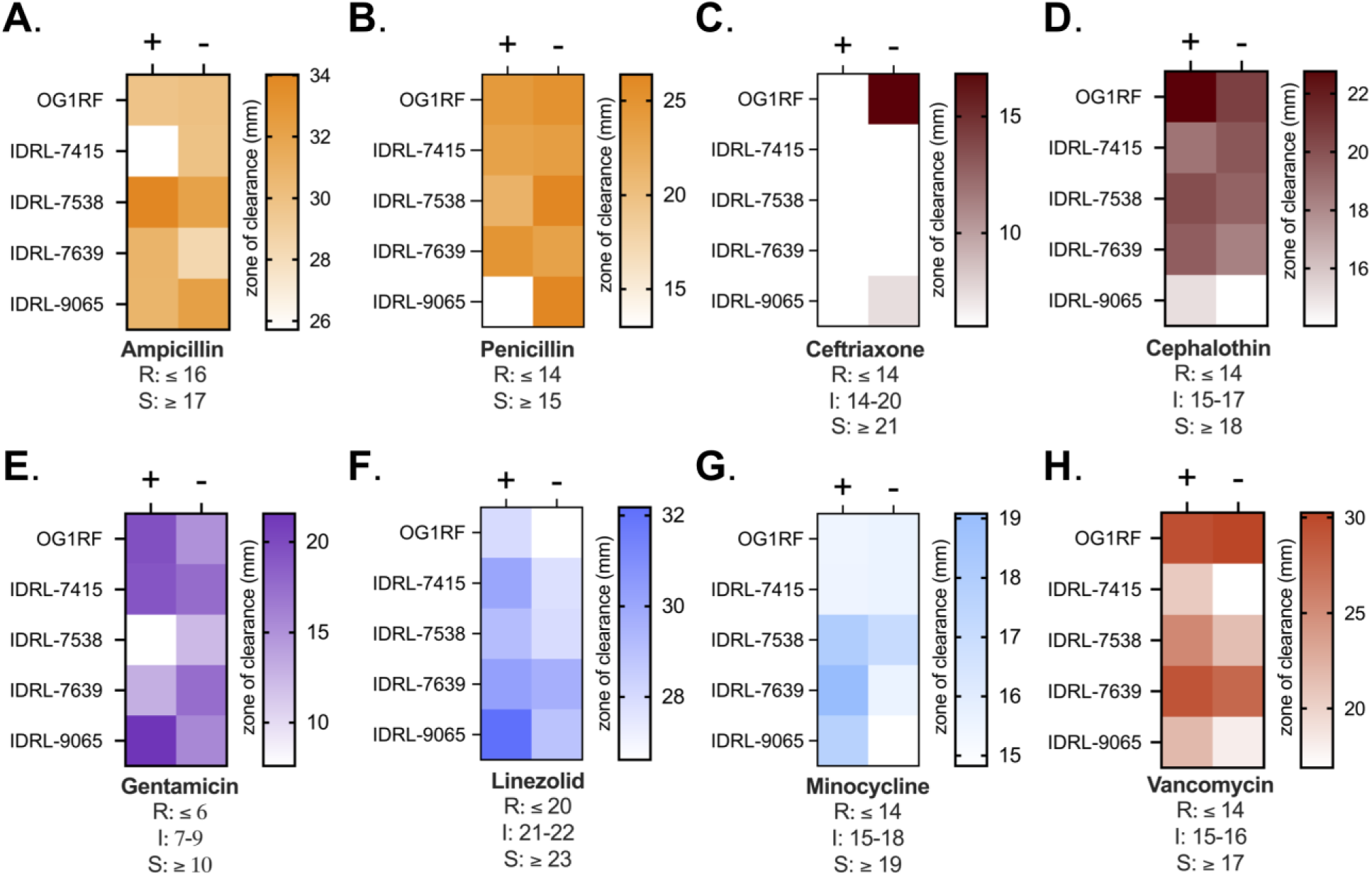
Susceptibility phenotypes of PJI isolates to antimicrobials. **(A-H)** Disk diffusion assays were used to observe PJI isolates response to different classes of clinically relevant antibiotics on agar plates supplemented with 30% SSF (+) or 30% H_2_O as a control (-). Data represent an average of 3 independent biological replicates. Color-coding indicates classes of related antibiotics (orange = β-lactams, maroon = cephalosporins, purple = aminoglycoside, blue = oxazolidenone, light blue = tetracycline, red = glycopeptide).

Previous reports demonstrated that *S. aureus* only aggregates in SSF under shear conditions and not after static growth^41^. This was intriguing considering these *E. faecalis* PJI isolates aggregated significantly in SSF when grown statically, so we assessed aggregation of these isolates grown in SSF with shaking. Interestingly, aggregation was isolate-specific. OG1RF and IDRL-7639 had a 6- to 8-fold decrease (respectively) in aggregation when grown in SSF with shaking as compared to static but still had a slight increase in aggregation in SSF (**Supplementary** Figure 5). Furthermore, the significant increase in aggregation was lost in IDRL-7415, IDRL-7538 and IDRL-9065 when grown with shaking. As with other bacterial pathogens, *E. faecalis* aggregates in SSF. However, the growth conditions in which *E. faecalis* aggregates differ from those of *S. aureus*, providing insight into *E. faecalis* pathogenesis during a PJI.

### Growth in SSF alters *E. faecalis* susceptibility to antibiotics

Rising antibiotic resistance is a major concern for both commensal and pathogenic bacteria. Antibiotic prophylaxis prior to resection surgery of a PJI is a measure taken to reduce the risk of recurrent infection making it imperative to monitor effective antibiotic regimens. Based on the finding that SSF impacts *E. faecalis* growth, biofilm formation, and aggregation^40,41^ (**Figure 4 and 5**), we decided to measure antibiotic susceptibility of these isolates when grown on agar supplemented with SSF using disk diffusion assays. We did not find any general changes in susceptibility across all strains, but instead observed strain-specific differences with individual antibiotics. IDRL-7415 was less susceptible to ampicillin and cephalothin but more susceptible to gentamicin, linezolid and minocycline when grown in SSF. Additionally, IDRL-9065 was less susceptible to β-lactam antibiotics but more susceptible to gentamicin, linezolid, minocycline and vancomycin. IDRL-7358 had increased resistance to gentamicin in SSF but was more susceptible to linezolid and minocycline. This is consistent with the presence of a gentamicin resistance gene in this isolate and our MIC assay results (**Table 4**). These findings suggest that antimicrobial efficacy against *E. faecalis* is altered when cells are grown in SSF. Further investigation into the mechanisms driving these changes in SSF may help guide improved antibiotic treatment regimens.

**Figure 5.**
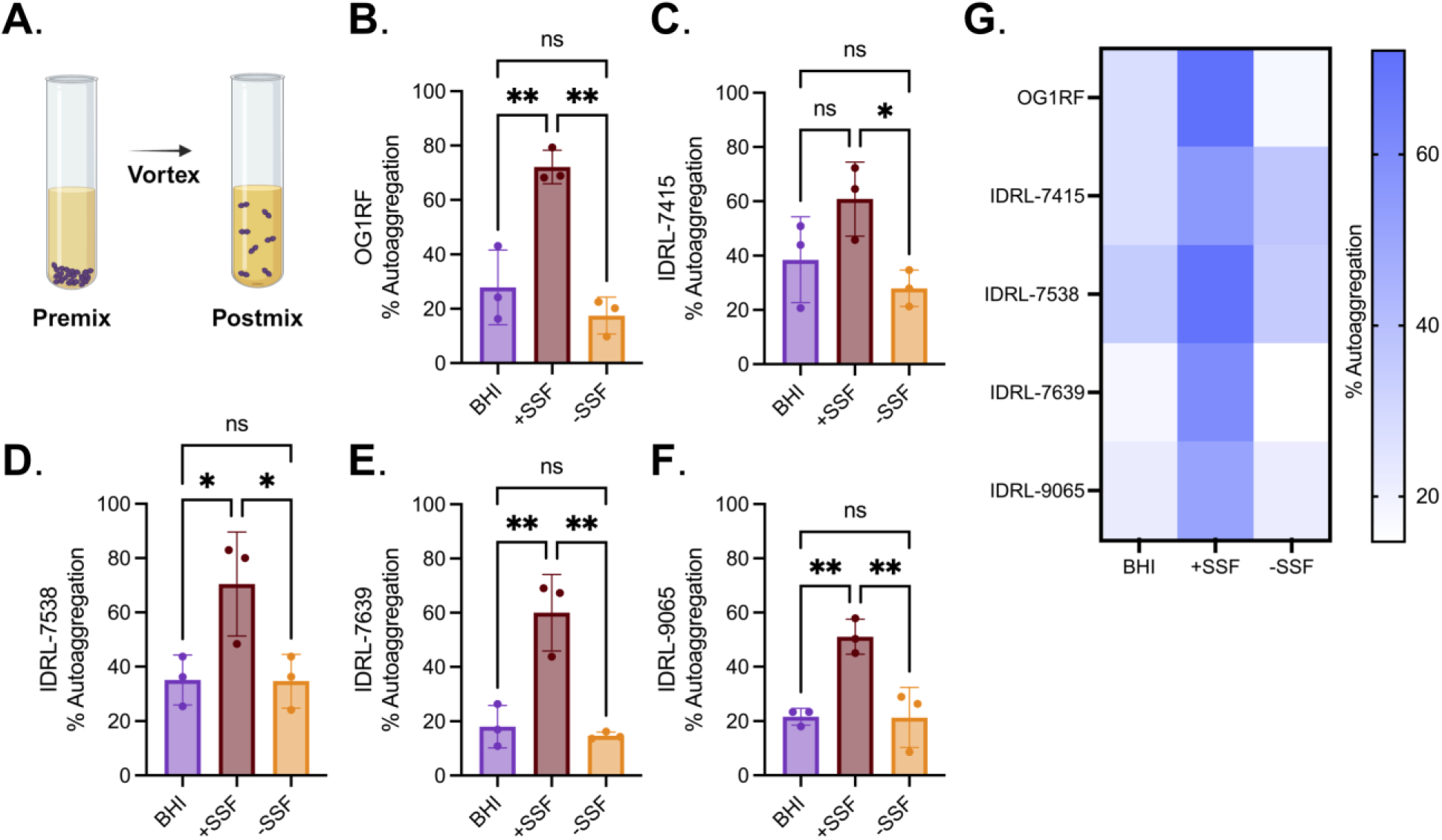
PJI isolates auto aggregation increases when grown in SSF conditions. **(A)** Schematic of autoaggregation protocol. Each clinical isolate was grown in 100% BHI (BHI), 70% BHI + 30% SSF (+SSF) and 70% BHI + 30% H_2_O (-SSF) for 16-18 h at 37°C, after which percent autoaggregation was calculated in each growth condition. Individual values are shown for **(B)** OG1RF, **(C)** IDRL-7415, **(D)** IDRL-7538, **(E)** IDRL-7639, **(F)** IDRL-9065. Summary data is visualized as a heat map in **(G)**, illustrating aggregation averages between each isolate in each condition. Statistical significance was determined using ordinary one-way ANOVA with Tukey’s multiple comparisons. ns = P > 0.05, * = P ≤ 0.05, ** = P ≤ 0.01, *** = P ≤ 0.001.

## Discussion

In this study, we show the first genomic and phenotypic analysis of *E. faecalis* isolated from periprosthetic joint infections (PJI). Previously, *E. faecalis* PJI investigations have been limited to retrospective cohort studies describing the prevalence and challenges associated with *E. faecalis* PJIs^12,45^. Therefore, particular emphasis was placed on analysis of factors relevant to *E. faecalis* infections, including biofilm formation and antibiotic resistance. Our genomic findings reveal differences in antibiotic resistance genes, virulence factors and prophages present in each genome. Plasmids, including some predicted to be pheromone-inducible conjugative plasmids, were found in three of the four strains. More investigations will be necessary to confirm this observation and determine whether plasmid transfer is critical for establishment and persistence of PJIs. We also identified complete prophages within three of the four strains, including two prophages previously identified in *E. faecalis* isolates from the oral cavity and bacteremia^35,36^. These results support previous findings describing genome diversity in *E. faecalis* clinical isolates^16,46^ and demonstrate that studying multiple strains of *E. faecalis* can provide insight into different traits associated with infection.

*E. faecalis* strains not only have diverse genomic features but also have differences in biofilm morphology^16^. The results reported here support these findings. IDRL-7415, the most genetically related isolate to OG1RF, had similar biofilm architecture and biofilm biomass accumulation relative to OG1RF. Conversely, isolates that were more distantly related to OG1RF, IDRL-7639 and IDRL-9065, had biofilm architectural differences such as chaining and aggregation. We also found variation in biofilm biomass, as determined by safranin staining, across the PJI isolates when grown in BHI. There was more variation during early biofilm growth (6 h) compared to late biofilm growth (24 h). Despite these changes in biofilm biomass and morphology, biofilms had similar viable cell counts at both 6 and 24 h. This suggests that these isolates may have differences in extracellular matrix material that contributes to overall biofilm biomass. This aligns with previous studies that showed strain-specific differences in matrix composition ^16,47,48^. Further investigation of biofilm matrix composition in *E. faecalis* PJI isolates could guide the development of anti-biofilm treatments.

Synovial fluid is a complex, viscous mix with a high abundance of hyaluronic acid^49^, albumin^40^, fibrinogen, and fibronectin^50^. Exposure to synovial fluid has been shown to affect growth, antibiotic resistance, biofilm matrix, and aggregation for other PJI pathogens, including *S. aureus*^37,38,46–48,51^. Our results show that *E. faecalis* grows less in SSF. However, it is unknown whether SSF reduces viability of *E. faecalis*, or if *E. faecalis* is unable to acquire necessary nutrients for survival in this environment. Future functional genomic investigations would provide insight into these mechanisms. We also found strain-specific differences between *E. faecalis* PJI isolates when grown in SSF, including changes in biofilm architecture and aggregation. These results suggest that *E. faecalis* biofilms develop differently when grown in SSF. However, growth in SSF did not result in significant differences in biofilm index, which takes into account safranin-stained biofilm material relative to overall cell growth. We also found that all four isolates and OG1RF had significantly increased aggregation during planktonic growth in SSF compared to the control condition. This suggests that SSF-mediated aggregation may not be unique to PJI isolates, but may be a general response of *E. faecalis* during growth in synovial fluid. It is important to note that we did not see a correlation between biofilm-forming capacity and SSF-induced aggregation. This suggests that these phenotypes may still be important, but independent of each other during PJI. However, it still remains unclear which components contribute to *E. faecalis* aggregation in SSF and whether aggregation is a temporary state during infection. This warrants further investigations into the kinetics of SSF-induced *E. faecalis* aggregation and the host components that mediate aggregation. The results described here are the first description of biofilm architectural differences in SSF and provide insight into the survival mechanisms used by *E. faecalis* during a PJI.

Surgical intervention followed by prolonged antibiotics is the most common treatment plan for patients diagnosed with a PJI. However, *E. faecalis* is intrinsically resistant to myriad antibiotics making these infections difficult to treat^45^. Prior to this work, no studies have reported the efficacy of antibiotics against *E. faecalis* in synovial fluid. Here, strain-specific differences in susceptibility to antibiotics were observed when *E. faecalis* isolates were grown in SSF compared to control conditions. This supports the broader idea that environmental conditions and media choice may impact antibiotic efficacy, which underscores the need to pursue susceptibility testing in conditions that closely resemble the host region^49–52^. In conclusion, our findings provide the first evidence of genetic heterogeneity among *E. faecalis* PJI isolates along with strain-specific differences in biofilm morphology and antibiotic resistance when grown in SSF. These results provide a platform for future studies to better understand *E. faecalis* PJI pathogenesis and treatment failure.

## Materials & Methods

### Collection of Isolates

This study was conducted under Mayo Clinic Institutional Review Board study 09-000808. Briefly, explanted prostheses from subjects undergoing arthroplasty resection due to PJI were placed in Ringer’s solution. Biofilms were removed using a previously published protocol for vortexing and sonication^53^. *E. faecalis* was cultured from resulting sonicate fluids and frozen in a microbank (Prolabs) at -80°C. The sonicate fluids were collected between March 2005 and September 2009.

### Bacterial strains and growth conditions

Bacterial isolates were maintained as freezer stocks at -80°C in 25% glycerol. Isolates were routinely grown in BHI (brain heart infusion, BD Difco) broth for overnight cultures for all experiments unless otherwise indicated. When required, agar was added to the growth medium at a final concentration of 1% (wt/vol). Simulated synovial fluid (SSF) was obtained from Biochemazone (product code BZ183) and supplemented with the indicated amount of BHI. Growth curves were used to determine target conditions for SSF experiments. Briefly, overnight cultures of each strain were diluted 1:100 in 96-well plates (Corning Co-Star 3595) in 200 μL of 100% BHI, 70% BHI + 30% sterile H_2_O, and 30% SSF + 70% BHI and grown statically at 37°C. Absorbance was measured at 600 nm every 30 minutes for 16 h in a Biotek Epoch2 plate reader.

### Whole genome sequencing and analysis

For short read Illumina sequencing, *E. faecalis* isolates were subcultured from the freezer onto sheep blood agar. Whole genome sequencing was performed using Illumina Nextera XT library preparation and the Illumina MiSeq instrument and chemistry. For long-read sequencing, overnight cultures were grown in BHI for 16-18 h, after which ∼10^9^ cells were pelleted and stored at -20°C. Pellets were shipped on dry ice to SeqCoast for DNA extraction and sequencing. Briefly, DNA was extracted using the MagMAX Microbiome Ultra Nucleic Acid Isolation kit with bead beating. Sequencing libraries were prepared using the Oxford Nanopore SQK-LSK114 native barcoding kit with Long Fragment Buffer. Libraries were sequenced on the GridION platform using a FLOW-MIN114 Spot-ON flow cell (R10) with a translocation speed of 400 bps. Base calling was performed on the GridION using the super-accurate base calling model. All genome analysis was done using the Bacterial and Viral Bioinformatics Resource Center (BV-BRC, version 3.33.16). Full genomes were assembled using the Genome Assembly *(B)* service with Unicycler (v0.4.8)^54^, with default settings unless otherwise specified. Quality of genome assemblies was determined by Bandage plot analysis^15^. Genomes were annotated with RASTtk^55^, using the Genome Annotation service in reference to *E. faecalis* OG1RF (ATCC 47077, genome ID 474186.54). Bacterial phylogenetic trees were generated using the Bacterial Genome Tree service with RAxML^37^ with default settings. Genome size, antibiotic resistance genes, virulence factors, and plasmid replicons were identified using the Comprehensive Genome Analysis (*B*) service provided by BV-BRC. Protein family comparisons were performed using Comparative Systems Service in BV-BRC. Putative plasmids were analyzed in PlasmidFinder^18^, and prophages were identified using assembled genomes in PHASTER^34^.

### Gelatinase assays

Gelatinase activity was assessed using previously described methods^20,56^. Briefly, overnight cultures of each strain were grown in BHI media and spot plated on agar plates made from tryptic soy broth without added dextrose (TSB-D, BD Difco) supplemented with 3% (wt/vol) gelatin. Plates were incubated overnight at 37°C and then moved to 4°C for 1 h prior to imaging. Plate photos were obtained using a ProteinSimple (Cell Biosciences) FluorChem FC3 imager. Strains were considered gelatinase positive if they developed a halo around colony growth and gelatinase negative if no halo was present.

### Biofilm assays

96-well plate biofilm assays were carried out as described previously^21,57^. Overnight cultures were grown in BHI media at 37°C for 16-18 h. Overnight cultures were diluted 1:100 in the indicated medium, and 200 µL was added to a 96-well plate (Corning Costar 3595). Plates were incubated in a humidified plastic container at 37°C for the indicated length of time. Cell growth was measured in a Biotek Epoch2 plate reader with the absorbance at 600nm (OD_600_). Plates were gently washed three times with ultrapure water using a Biotek plate washer, dried in a biosafety cabinet overnight, and stained with 100 µL of 0.1% safranin (Sigma). Stained plates were washed three times and dried. The A_450_ was measured to quantify safranin-stained biofilm biomass. Biofilm production was evaluated as the ratio of stained biofilm biomass to overall growth (OD_450_/OD_600_).

Submerged Aclar biofilm assays were performed as previously described.^19^ Overnight cultures were adjusted to 10^7^ CFU/mL in BHI, and 1 mL of each isolate was added to a 24-well plate (Costar 3524) with a sterile 5 mm Aclar disc. Plates were incubated at 37°C in a tabletop shaker incubator (Labnet 311DS) at 100 rpm. After 6 h, planktonic cultures were transferred to a microcentrifuge tube. Aclar discs were washed by gentle shaking in KPBS and transferred to a microcentrifuge tube with 1mL KPBS (1 Aclar disc/tube). Tubes with planktonic cultures and Aclar discs were vortexed at 2,500 rpm for 5 min in a BenchMixer multitube vortexer (Benchmark Scientific) and then diluted (10-fold serial dilutions) in KPBS and plated on BHI agar plates to enumerate colonies. At least three biological replicates (each with two technical replicates) were performed for all strains.

### Fluorescence microscopy

For all experiments, 96-well plate biofilm assays were prepared as described above in triplicate replicates. Cultures were grown in black clear-bottom 96-well plates (Thermo Scientific Nunc 165305) in the indicated medium. Planktonic cells were removed with a multi-channel pipette, and biofilms were gently rinsed in KPBS three times. Biofilms were fixed with 100 µL formalin overnight at 4°C. After fixing, plates were gently rinsed in KPBS three times and stained with Hoechst 33342 for 15 min at room temperature.

### Microscopy and image processing

Images were captured using the Keyence BZ-X810 microscope with a Chroma DAPI filter (AT350/50x). For each technical replicate, two representative images were obtained, yielding 6 images per strain per condition from which a final representative image was chosen. Representative images were processed using the Fiji ImageJ package (version 2.9.0/1.53t) and subjected to background subtraction with a rolling-ball radius of 90 pixels using the internal ImageJ function as well as uniformly applied brightness and contrast adjustments of the entire image prior to cropping. Biofilms of each image were false colored with cyan, cropped to 500 by 500 pixels, and exported as PNG files.

### Autoaggregation assay

Each strain was inoculated into culture tubes containing 3 mL of indicated medium and grown statically for 24 h at 37°C. For each strain and condition, culture tubes were gently removed from the incubator (without disrupting any aggregation). Collection of samples was done via pipetting from the middle region of the undisrupted culture tube followed by a 1:5 dilution in a cuvette and measurement of absorbance at 600nm (A_600_). These were considered the pre-mixed culture values. Following this, a second measurement of the same strain and condition was performed by vortexing the culture tube for ∼3 seconds, sampling of the culture by pipetting at the middle of the tube, dilution (1:5) in a cuvette, and measurement of the A_600_. This was considered the post-mixed culture value, or the actual A_600_ of the culture. Autoaggregation was quantified as ((postmix - premix)/postmix)*100 to calculate the final percentage of aggregation in the culture.

To perform the shaking autoaggregation assay, each strain was inoculated into 3 mL of the respective medium listed above, then incubated at 37°C at 220 rpm on an Innova 2300 shaker (New Brunswick Scientific) for 24 h. Upon removal of the culture tubes from the shaker, the cultures were allowed to settle on the benchtop for 10 minutes undisturbed. Following this settling period, the cultures were subject to the same A_600_ measurement described above for the static cultures.

### Antibiotic susceptibility testing

Overnight cultures were grown in BHI for 16-18 h and plated on 1% BHI-agar plates supplemented with 30% sterile H_2_O or 30% SSF and allowed to dry. BD BBZ Sensei-Discs were placed on dried agar plates and incubated, face up, for 16-18 h. Antibiotics included ampicillin (10 µg), penicillin (10IU/IE/UI), ceftriaxone (30 µg), cephalothin (30 µg), gentamicin (120 µg), linezolid (30 µg), minocycline (30 µg) and vancomycin (30 µg). Agar plates were imaged on a ProteinSimple (Cell Biosciences) FluorChem FC3 imager and zones of clearance were measured using ImageJ package (version 2.9.0/1.53t). Three independent biological replicates were performed with freshly made plates.

### Statistical analysis

Replicates (technical and biological) and statistical tests are described in each figure legend. Statistical analysis was performed using GraphPad Prism (Version 10.0.3).

## Data availability statement

Files from Illumna and Nanopore sequencing and genome files have been deposited as BioProject PRJNA1069998.

## Acknowledgements

This work was supported by the National Institutes of Health (award number R01 AR056647 to RP and T32 GM140936 to ALH). The content is solely the responsibility of the authors and does not necessarily represent the official views of the National Institutes of Health. Additional support was provided by start-up funds from the University of Minnesota to JLEW. This work was also supported by the resources and staff at the University of Minnesota Genomics Center (RRID: SCR_012413). We thank members of the Willett lab for their helpful feedback on the manuscript.

Dr. Patel reports grants from MicuRx Pharmaceuticals and BIOFIRE. Dr. Patel is a consultant to PhAST, Day Zero Diagnostics, Abbott Laboratories, Sysmex, DEEPULL DIAGNOSTICS, S.L., Netflix and CARB-X. In addition, Dr. Patel has a patent on *Bordetella pertussis/parapertussis* PCR issued, a patent on a device/method for sonication with royalties paid by Samsung to Mayo Clinic, and a patent on an anti-biofilm substance issued. Dr. Patel receives honoraria from Up-to-Date and the Infectious Diseases Board Review Course.

